# Extinction burst could be explained by curiosity-driven reinforcement learning

**DOI:** 10.1101/2024.08.28.610088

**Authors:** Kota Yamada, Hiroshi Matsui, Koji Toda

## Abstract

Curiosity encourages agents to explore their environment, leading to learning opportunities. Although psychology and neurobiology have tackled how external rewards control behavior, how intrinsic factors control behavior remains unclear. An extinction burst is a behavioral phenomenon in which a sudden increase in the frequency of a behavior immediately follows the omission of a reward. Although the extinction burst is textbook knowledge in psychology, there is little empirical evidence of it in experimental situations. In this study, we show that the extinction burst can be explained by curiosity by combining computational modeling of behavior and empirical demonstrations in mice. First, we built a reinforcement learning model incorporating curiosity, defined as expected reward prediction errors, and the model additively controlled the agent’s behavior to the primary reward. Simulations revealed that the curiosity-driven reinforcement learning model produced an extinction burst and burst intensity depended on the reward probability. Second, we established a behavioral procedure that captured extinction bursts in an experimental setup using mice. We conducted an operant conditioning task with head-fixed mice, in which the reward followed after pressing a lever at a given probability. After the training sessions, we occasionally withheld the reward delivery when the mice performed the task. We found that phasic bursts of responses occurred immediately after reward omission when responses were rewarded with a high probability, suggesting that the magnitude of reward prediction errors controlled the burst. These results provide theoretical and experimental evidence that intrinsic factors control behavior in adapting to an ever-changing environment.

**Significance statement:** In theories of learning and behavior, primary rewards such as food, water, and sex occupy a dominant position as factors controlling behavior. However, primary rewards are scarce. Experimental investigations in psychology, neuroscience, ethology, and economics have revealed that novelty, uncertainty, and unpredictability drive behavior. How these intrinsic factors affect behavior is essential for a comprehensive understanding of the principles of learning and behavior. This study provides theoretical and experimental evidence that operant responses in mice are directly controlled by external rewards and intrinsic factors such as curiosity. Our study provides a robust example of curiosity-driven behavior and paves the way for understanding the mechanism of curiosity.

## Introduction

In an ever-changing environment, organisms must maximize the rewards of their actions. Although primary rewards, such as food and water, robustly control behavior, the world is too vast to earn all possible rewards. In such environments, animals have few opportunities to learn actions that directly lead to rewards. To overcome this problem, both animals and artificial agents must expand their learning opportunities by exploring vast and uncertain environments.

Artificial agents also face difficulties in learning sets of actions that maximize rewards in an environment where rewards are sparsely distributed (1). Efforts have been made to resolve this problem by additively incorporating curiosity into external rewards to encourage agents to explore their environment actively (2–6). Although curiosity can be formulated in several ways, they assume that agents obtain information when observing an event. One typical formulation is based on prediction errors (2, 3, 6). If an agent is familiar with an event, the agent can produce the correct prediction. In other words, if an event produces a large prediction error, the event is a novel situation or a change in the environment for the agent, implying that opportunities for learning remain. Thus, when curiosity is defined based on prediction errors, it encourages the agent to explore new or unpredictable states in the environment.

The pursuit of uncertainty, novelty, and unpredictability in animals is known to facilitate learning. Animals also display orientation, preference, and exploration of novel, unpredictable, and informative events and situations (7–15). It has long been reported that a novel stimulus for an individual induces a orienting response of the individual (11, 16, 17) and animals exhibit an orienting response to stimuli that produce prediction errors (16, 18); such errors also promote learning by modulating parameters associated with learning (19–22). When the outcome of a response deviates from the prediction or is filled with uncertainty, animals increase their exploration tendencies or change their behavioral strategies (23–26). This behavioral evidence demonstrates that intrinsic factors expand learning opportunities and promote learning (27, 28).

When an animal faces unpredictable reward omission for an operant response, responses occur more frequently immediately after omission. This phenomenon is known as an extinction burst (29). For example, when we press a key on a laptop and no character appears on the display, we sometimes press the key repeatedly. Although the extinction burst is referred to anecdotally, it is reported to occur when stopping problematic behavior using extinction in clinical situations (30, 31), and seems to be a familiar phenomenon in everyday life, there is little experimental evidence showing of it (32, 33). Extinction produces several byproducts. It induces responses other than the target operant response, such as aggression, social interaction, and drinking behavior (34–36) and increases the variability of the target response (26). Because most existing studies on extinction in the context of operant conditioning have been conducted in a conventional operant box, in which animals behave freely, the side effects of extinction might spread across a broad repertoire of phylogenetically inherited behavior, making it difficult to observe the extinction burst in the experimental situation. Recently, neuroscience has begun to use a head-fixed apparatus that limits the spontaneous behavior of mice, examines target behavior, and records physiological activities effectively (37). Utilizing this head-fixed procedure in mice, we predicted that minimizing experiment-unrelated behaviors might be efficient in demonstrating the extinction burst.

In addition, standard reinforcement learning and other conventional learning theories cannot explain extinction bursts because the number of responses increases phasically, although the response is not rewarded (38–40). Thus, it remains unclear whether the extinction burst actually exists and what factors determine it. Despite the lack of empirical evidence, the extinction burst might be explained in terms of curiosity because animals experience reward prediction errors when a reward is unpredictably withheld.

We thus provide theoretical and experimental evidence that curiosity controls behavior through computational simulations using a curiosity-driven reinforcement learning model and experimental investigations of operant conditioning experiments with mice. Three main efforts have been made to achieve this goal. First, we built a curiosity-driven reinforcement learning model incorporating retrospective curiosity, defined as the expected reward prediction error (20), which controls the frequency of the response in parallel with the primary reward. Second, we conducted computational simulations using curiosity-driven and standard reinforcement learning models to clarify whether the curiosity-driven model could predict the behavioral phenomenon of the extinction burst. We also conducted simulations under several parameter settings to identify the control variables for the extinction burst. Third, we conducted an operant conditioning task to examine whether the identified control variables observed in the computational model explained the extinction bursts observed in the experimental data.

## Results

### Q-learning with Hierarchical Curiosity Module

We built a curiosity-driven reinforcement learning model called Q-learning with Hierarchical Curiosity Module (Q-HCM) that incorporates curiosity, defined as the expected reward prediction error, as per Fig. 1A; (20). This model uses a vanilla Q-learning model (VQM) as its backbone. VQM estimates action-value function *Q*(*s_t_, a_t_*) which represents the expected value of the reward acquired by an action *a*_*t*_ in a given state *s*_*t*_ using the delta rule. There is only one state and action in our experiment (see Materials and Methods for details); therefore, we denote *Q*(*s*_*t*_, *a*_*t*_) as *Q*. *Q* is updated with the reward prediction error when an agent emits a response based on whether a reward was acquired by the response. Reward prediction error, denoted as δ_*t*_, is the difference between the predicted reward by action *a*_*t*_ and observed reward r_*t*_:

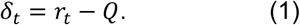

**Figure 1.**
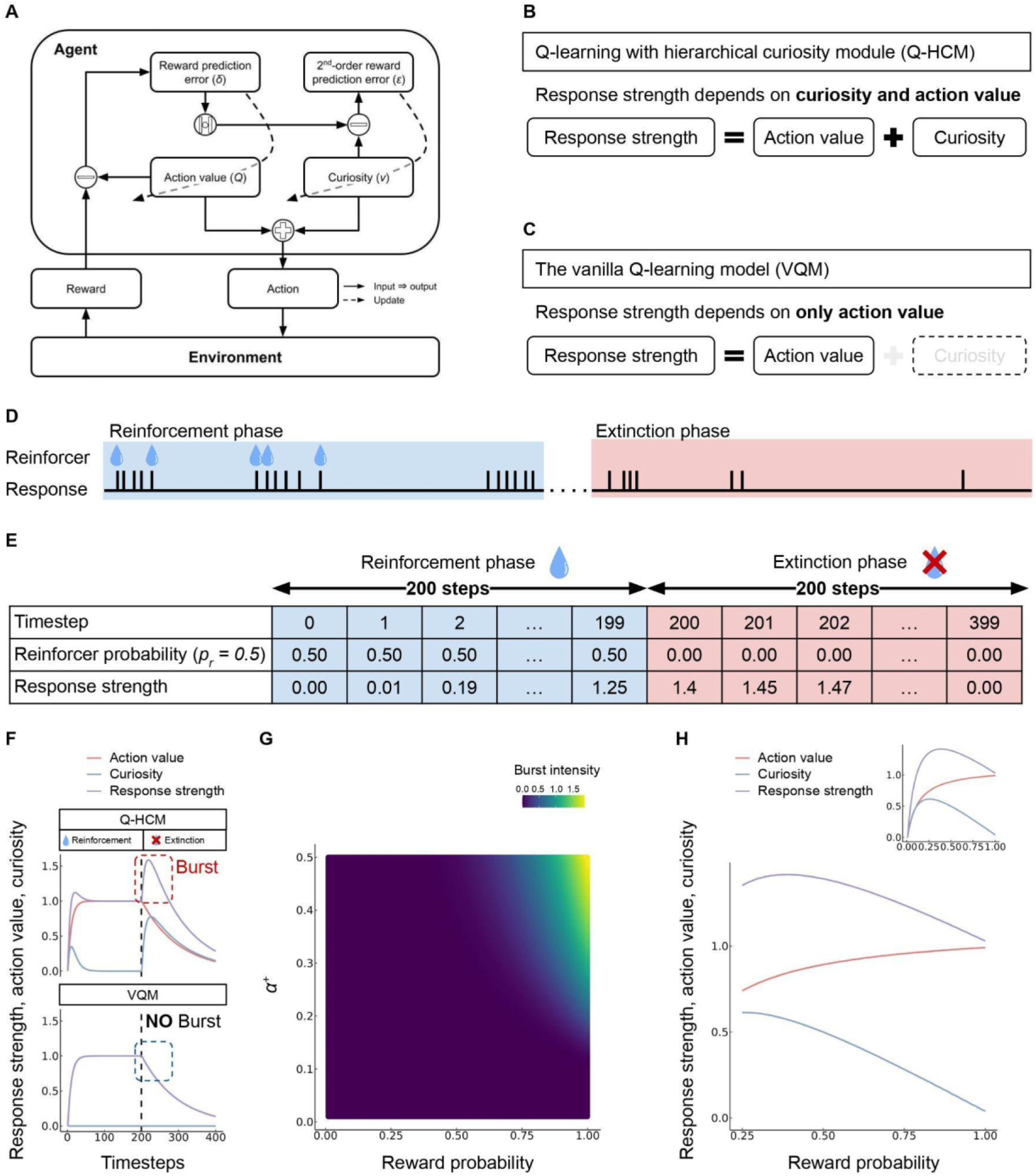
Simulation results. **A.** Schematic representation of Q-learning with Hierarchical Curiosity Module (Q-HCM) **B.** Simplified schematic representation of Q-HCM where both action value and curiosity determine the response strength. **C.** Simplified schematic representation of VQM, where only action value determines the response strength. **D.** Example of raster plots of response and reward in a VR 4 schedule and extinction. **E.** Example of simulations. We calculate the expected response strength at each timestep under VR 1 (*p*_r_ *=* 1.0) with the given parameters (α^+^ = *0*.*1*, α^−^ = *0*.*01*, α^*v*^ = *0*.*1*). Here, response strength is defined as the sum of curiosity and action value without the weight parameter. **F.** Examples of temporal changes in response strength, action value, and curiosity generated by Q-HCM (top) and VQM (bottom). The red, blue, and purple lines in the upper panel indicate action value, curiosity, and response strength, respectively,. Response strength is the same as action value in VQM, and each color indicates the response strength generated by different parameter sets of the VQM. The first half 200 steps are the reward phase and the last half steps are the extinction phase in each panel. **G.** The heatmap shows the relationship between burst intensity, and combination of learning rate and reward probability. The brighter colors indicate larger burst intensity that is the area outlined by the red dashed area in **F**. **H.** Relationship between the response strength and the reward probability. The large panel shows the range from 0.25 to 1.00 where we employed the mice’s experiments and the upper right panel shows the full range of reward probabilities. The red, blue, and purple lines indicate the action value, curiosity, and response strength, respectively. The model parameters were α^+^ = *0*.*1*, α^−^ = *0*.*01*, α^*v*^ = *0*.*1*.

When the observed reward is not the same as the action value, *Q*, it is updated by equation *Q* ← *Q* +αδ_*t*_. α denotes the learning rate which determines how much the agent updates the action value at one step. This is the basic form of Q-learning. We assume different learning rates for the positive and negative prediction errors. The asymmetric update rule is as follows:

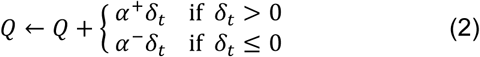

Next, we extend the VQM to Q-HCM. The primary extension is that curiosity additively controls the response to the action value. We define curiosity as the predicted reward prediction error obtained by integrating the second-order reward prediction error over time. The second-order reward prediction error, denoted as ϵ_*t*_, is difference between predicted reward prediction error, denoted as *v*, and the absolute value of actual reward prediction error |δ_*t*_|:

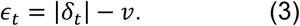

Then the predicted reward prediction error*v* is updated using the following equation:

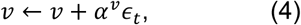

where *v* denotes how much the action *a*_*t*_ causes a reward prediction error. Finally, curiosity controls the response in parallel to the action value:

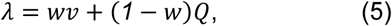

where *w* is a weighting parameter for *Q* and *v*, and its range is limited 0 to 1. When *w* = 0, the response is determined solely by action value *Q* and Q-HCM is reduced to VQM.

### Curiosity is essential for generating the extinction burst

We compared the behaviors of two different models, Q-HCM and VQM, during extinction to confirm that curiosity is necessary for extinction bursts. In Q-HCM, the response strength is determined by the action value and curiosity (Fig. 1B). By contrast, response strength depends only on the action value in the VQM (Fig. 1C). The simulated environment contains two phases (Fig. 1D). In the reward phase, agents can obtain a reward for each action at a given probability, termed the variable ratio (VR) schedule. The extinction phase, in which the agents cannot obtain any reward regardless of their actions, follows the reward phase. Because reward presentation depends only on the agents’ actions in the VR schedule, the passage of time has no effect on reward probability. This time-independence allowed us to simplify the simulation environment. We thus only calculated the action value and curiosity at each time step based on whether the reward occurred in the reward phase (Fig. 1E; see the detailed simulation procedure in Materials and Methods). The reward phase continued for 200 time steps, and the response strength reached an asymptote in both models (Fig. 1F). The extinction phase followed immediately after the reward phase. While action values decayed monotonically in both models, curiosity and response strength increased phasically in Q-HCM (Fig. 1F), suggesting that curiosity was necessary for the extinction burst.

We conducted further simulations and analyses of the Q-HCM behavior to identify what controls the extinction burst. We simulated the behavior of the Q-HCM under several parameter settings and calculated the burst intensity for each setting. We defined the burst intensity as the increase in response strength after extinction (the area outlined by the red dashed area in Fig. 1F). To correctly calculate burst intensity, we calculated the response strength until it reached an asymptote during the reward phase. We then introduced an extinction phase, which continued until the response strength became lower than that of the asymptote in the reward phase. Burst intensity was higher when the reward probability and learning rate were higher (Fig. 1G), suggesting that reward probability was the primary factor controlling the extinction burst.

Q-HCM displayed unique behavior not only in the extinction phase but also in the reward phase. Although the action value increased monotonically as the reward probability increased, curiosity and response strength decreased at the highest reward probabilities (Fig. 1H). Because VQM lacks curiosity and the response strength depends on the action value, it shows a monotonic relationship between reward probability and response strength, suggesting that the downturn in the response rate at the highest reward probability is unique for Q-HCM.

### Extinction burst was notable under high reward probability in the lever pressings of head-fixed mice

The simulations showed that curiosity caused an extinction burst and that the reward probability determined burst intensity. Here, we examined whether the mice showed an extinction burst, which is more likely to occur in a high reward probability condition. We employed the operant conditioning task with VR schedules, where the reward could be obtained by lever pressing at a given probability (Fig. 1D and 2A; 41). There were four reward probabilities (1.0, 0.5, 0.33, and 0.25 for VR 1, 2, 3, and 4, respectively). Two hundred rewards were delivered in one session, and we trained the mice under each condition until they completed 200 reward in five consecutive days. After the training, extinction trials were introduced during the sessions. Extinction trials were inserted without any cue that signaled the start of the extinction trial during the sessions, four times randomly (Fig. 1B), and continued until the mice stopped pressing the lever for 1 min. In the test session, the mice pressed the lever and obtained rewards and stopped pressing the lever after the onset of extinction (Fig. 2C). We conducted four extinction sessions per condition, and all mice experienced all four reward probability conditions.

**Figure 2.**
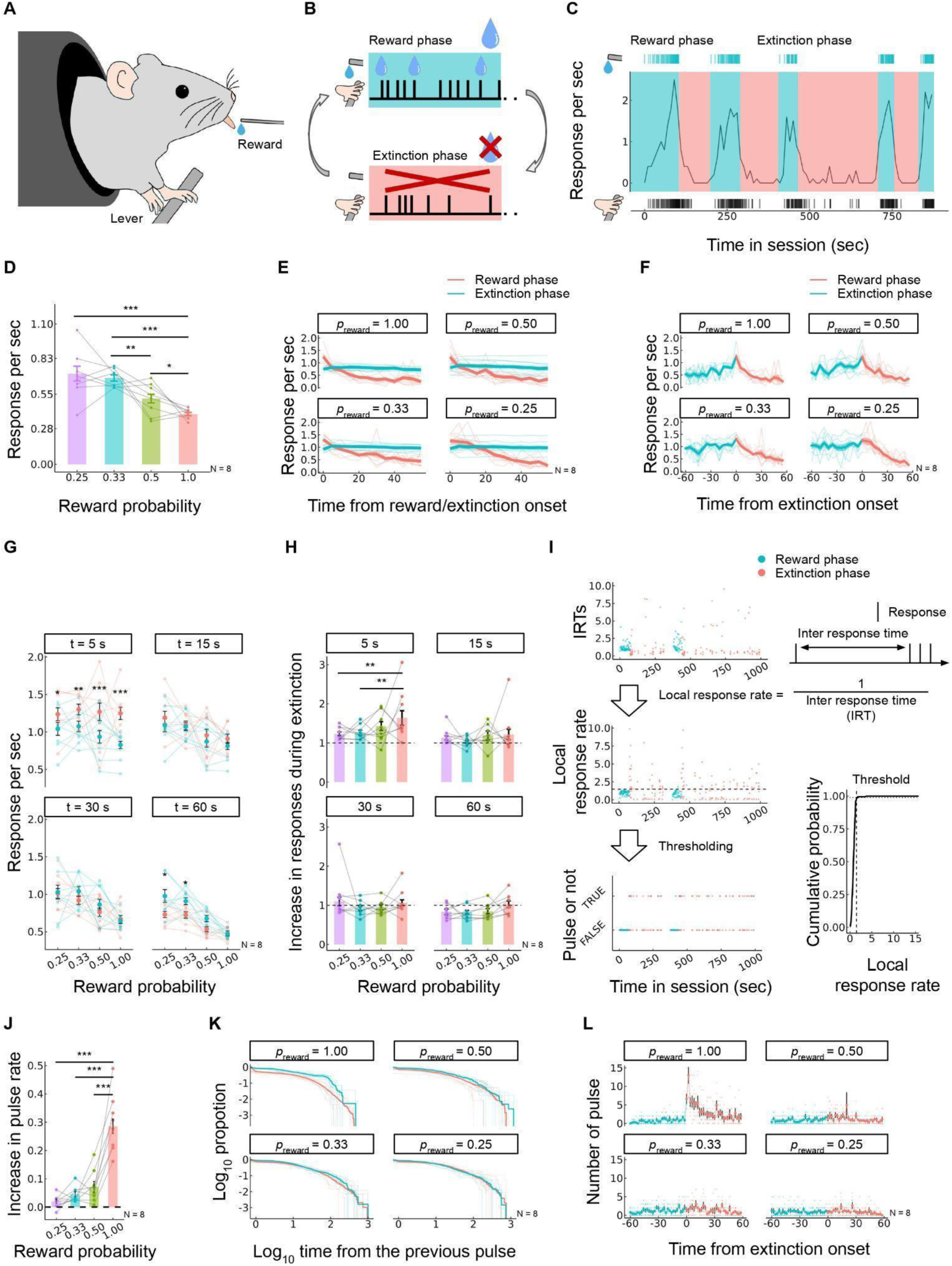
Results of the behavioral task with head-fixed mice. A. Schematic representation of the experimental setup for operant conditioning with head-fixed mice. B. Overview of the test session including the reward phase and the extinction phase. The reward and the extinction phases were conducted alternatively during the session. C. An example of raster plots of a mouse’s lever pressing (vertical black bars in bottom), rewards (vertical blue bars in top), and average number of responses in 5-second bins. D. The relationship between response rates of lever pressing during the reward phase and the reward probabilities. The points indicate the data from individual subjects. E. Temporal changes in response rates from onsets of the reward (red lines) and extinction (blue lines). F. Temporal changes in response rates around the onsets of extinction. G. Response rates around the onsets of extinction with specific time windows (5s, 15s, 30s, and 60s). H. Ratio of response rates during the extinction phase to those of the reward phase. I. Example of pulse detection analysis (see Materials and Methods for details). J. Ratio of pulse rate during the extinction phase to the reward phase. K. Survival function of inter-pulse intervals during the reward phase (blue) and extinction phase (red). L. Number of pulses 60 seconds before and after the onset of extinction averaged across all subjects and sessions. The error bars indicate standard error of the mean.

First, we examined the relationship between reward probabilities and response rates to validate the Q-HCM prediction of a downturn in the response rate at the highest reward probabilities (Fig. 1H). In the highest reward probability condition, the response rate was the lowest of all four conditions (Fig. 2D), corresponding to the Q-HCM prediction. The linear-mixed model also revealed significant effects of the reward probability on the response rate (*F*_(3, 63.72)_ = 20.90, *p* < 0.001) and the response rates were lower for higher reward probabilities than lower reward probability (1.00 vs. 0.50, *t*_(63.84)_ = –2.68, *p* = 0.046; 1.00 vs. 0.33, *t*_(63.52)_ = –6.34, *p* < 0.001; 1.00 vs. 0.25, *t*_(63.18)_ = –6.75, *p* < 0.001; 0.50 vs. 0.33, *t*_(61.91)_ = –3.72, *p* = 0.002; 0.50 vs. 0.25, *t*_(64.46)_ = –4.22, *p* < 0.001; 0.33 vs. 0.25, *t*_(63.79)_ = –0.72, *p* = 0.89). The downturn in the response rate at the highest reward probabilities is a unique behavior of the Q-HCM; thus, this result supports the idea that the mice tracked reward uncertainty and controlled their behavior based on uncertainty.

We examined another prediction of Q-HCM, in which the reward probability was the primary controlling factor of the extinction burst (Fig. 1G). In the reward phase, response rates stabilized over time (blue lines in Fig. 2E and F). By contrast, response rates decreased with time during the extinction phase (red lines in Fig. 2E and F). However, response rates were higher in the reward phase immediately after extinction onset. To examine whether response rates increased immediately after extinction onset, we compared the response rates at 5, 15, 30, and 60 s before and after extinction onset. Response rates were larger in the extinction phase than in the reward phase with a 5-s time window analysis, but were not different with a longer time window analysis (Fig. 2G). The linear mixed model revealed a significant interaction between the time window and phase (*F*_(3, 624.96)_ = 17.93, *p* < 0.001) and the effects of reward probability (*F*_(3, 626.39)_ = 35.41, *p* < 0.001). Multiple comparisons supported that the response rates were higher in the extinction phase than the reward phase in the shortest time window (Reward vs. extinction with 1.00, *t*_(625.00)_ = 4.75, *p* < 0.001; Reward vs. extinction with 0.50, *t*_(625.00)_ = 3.84, *p* < 0.001; Reward vs. extinction with 0.33, *t*_(625.00)_ = 2.61, *p* = 0.009; Reward vs. extinction with 0.25, *t*_(625.00)_ = 2.03, *p* = 0.041) but lower in the longest time window with lower reward probabilities (Reward vs. extinction with 1.00, *t*_(625.00)_ = 0.06, *p* = 0.95; Reward vs. extinction with 0.50, *t*_(625.00)_ = –1.64, *p* = 0.101; Reward vs. extinction with 0.33, *t*_(625.00)_ = –2.12, *p* = 0.034; Reward vs. extinction with 0.25, *t*_(625.00)_ = –2.58, *p* = 0.010). The increase in response rates at the onset of extinction was larger in the highest reward probability, and decreased as the reward probability decreased; it also depended on the time window (Fig 2H). The linear mixed model revealed significant effects of reward probability and time window (reward probability: *F*_(3, 306.82)_ = 5.26, *p* < 0.001; time window: *F*_(3, 296.47)_ = 24.42, *p* < 0.001). Multiple comparisons also revealed that the difference was larger than the highest reward probabilities and lower reward probability in the shortest time window (1.00 vs. 0.33, *t*_(300.74)_ = 3.17, *p* = 0.009; 1.00 vs. 0.25, *t*_(300.40)_ = 3.25, *p* = 0.007).

As the extinction burst is described as a phasic burst of response, we detected and analyzed unusual response speeds. We defined the local response rates as the reciprocal of the inter-response times (IRTs) for each response (Fig. 2I, top and middle left-hand panels). We classified each response into a pulse based on whether the local response rate was over the 95% quantile of the local response rates during the reward phase for each session and subject (Fig. 2I, middle right and bottom panels). We compared the pulse rates during the reward and extinction phases, and found that the pulse rate increased in the extinction phase only under the highest reward probability condition (Fig. 2J). The linear mixed model revealed significant effects of the reward probability (F_(3, 72.76)_ = 54.07, *p* < 0.001) and the pulse rates were larger in the highest reward probability than others (1.00 vs. 0.50, *t*_(64.27)_ = 9.22, *p* < 0.001; 1.00 vs. 0.33, *t*_(63.71)_ = 10.40, *p* < 0.001; 1.00 vs. 0.25, *t*_(64.29)_ = 11.02, *p* < 0.001). The inter-pulse intervals were shorter in the highest reward probability condition (Fig. 2K). Pulses occurred frequently at the onset of extinction (Fig. 2L). Considering these results, mice lever pressing was considered to be under the control of both primary rewards and reward uncertainty.

### Q-HCM model captures mice’s lever pressings under extinction

Experimental results for mice showed that phasic unusually fast responses, pulses, occurred with extinction. We compared a model incorporating curiosity (Q-HCM) and a model lacking curiosity (VQM) fitted to the empirical data to confirm that the burst of pulses was driven by curiosity. Because the extinction burst was transient (Fig. 2G, H, and L) and burst-like responses occupied limited parts of the overall data, we fitted the model to the dynamics of the pulse instead of the individual response. We assumed that the probability of pulse occurrence was mediated by action value and curiosity, and estimated the changes in action value, curiosity, and pulse occurrence probability over time based on the actual response, whether that response was a pulse, and the presence of a reward. For each subject’s session-by-session data, we estimated a set of parameters for the models that would fit the pulse time series. We calculated the WAIC to compare the fits of the two models to the empirical data. Under the highest reward probability condition, the WAIC of Q-HCM had a WAIC smaller than that of the VQM (Fig. 3A). The difference in the WAIC between Q-HCM and VQM decreased as the reward probability decreased (Fig. 3A). The increases in pulse occurrence probabilities with extinction matched the empirical and predicted data (Fig. 3B; *t*_(30)_=19.667, *p* < 0.001), indicating that Q-HCM successfully captured the behaviors of individuals.

**Figure 3.**
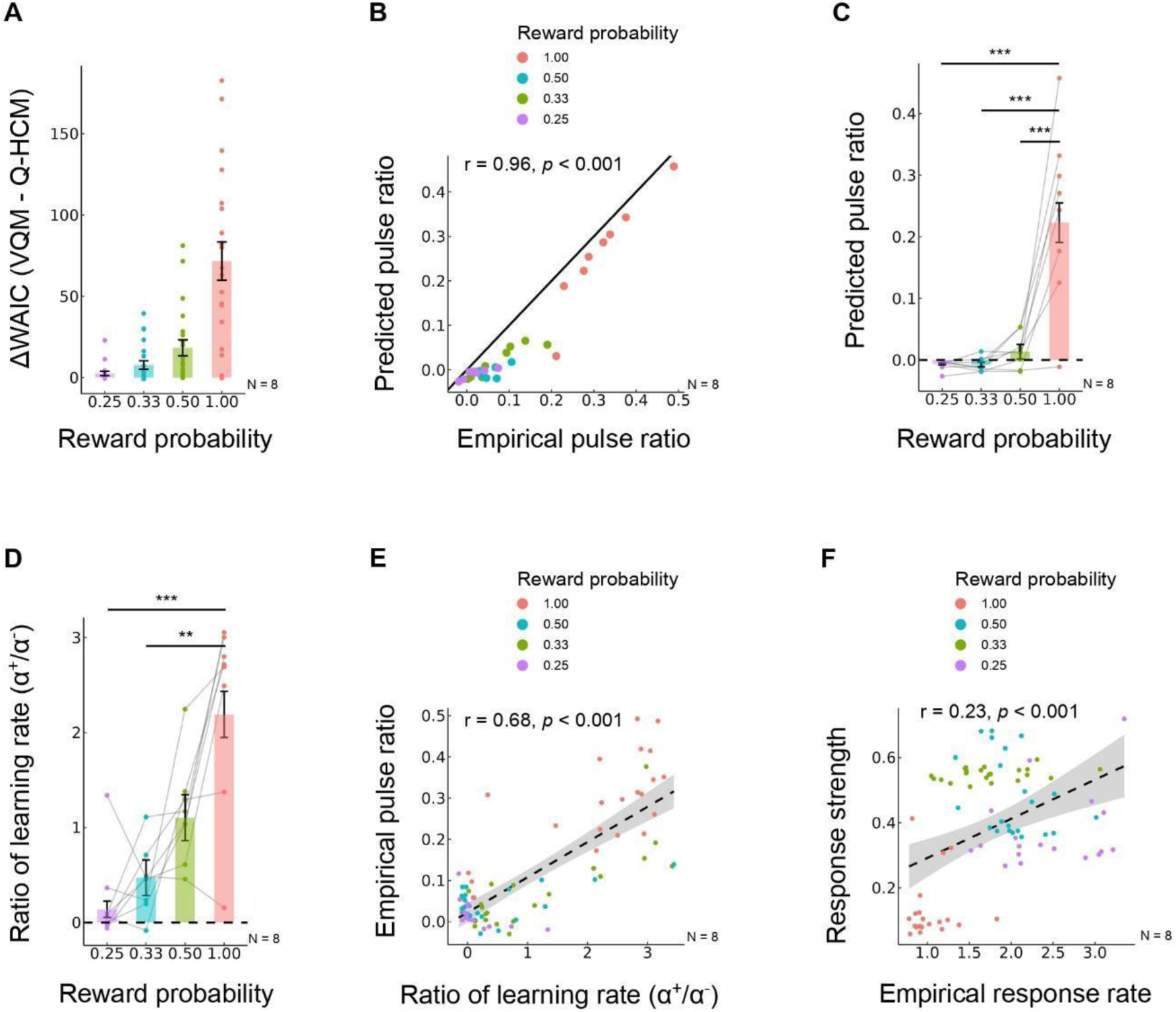
Results of the model fitting to the empirical data. A. Difference of WAIC between VQM and Q-HCM (VQM – Q-HCM) for each subject and session. If the difference is larger, Q-HCM is more well-fitted for the empirical data. B. The relationship between empirical pulse ratio and pulse ratio predicted by Q-HCM. The pulse ratio represents the ratio of the proportion of pulse in the reward phase to that during the extinction phase. Each point is averaged across sessions for each subject and condition. The point colors indicate reward probability in the reward phase. C. Predicted pulse ratio for each condition. D. Ratio of learning rate for positive reward prediction errors and that of negative prediction errors for all conditions. E. The relationship between ratio of learning rate and empirical pulse ratio. Each point represents data per session for an individual subject. The point colors indicate the reward probability in the reward phase. F. The relationship between empirical response rate during the reward phase and response strength estimated by the Q-HCM during the reward phase. The response strength is the weighted sum of action value and curiosity (eq. 5). Error bars indicate standard error of the mean.

Q-HCM reproduced the experimental results in which the burst responses occurred almost exclusively under extinction. Pulses were more likely to occur during the extinction phase in the higher reward probability condition (Fig. 2G), and the model showed similar results (Fig. 3C). The linear mixed model revealed significant effects of the reward probability (*F*_(3, 72.66)_ = 49.40, *p* < 0.001), and the predicted pulse ratio was larger for the highest reward probability than for the others (1.00 vs. 0.50, *t*_(69.80)_ = 9.19, *p* < 0.001; 1.00 vs. 0.33, *t*_(69.42)_ = 10.18, *p* < 0.001; 1.00 vs. 0.25, *t*_(68.90)_ = 9.79, *p* < 0.001). The simulations predicted that extinction bursts were more likely to occur when the learning rate for positive prediction errors was higher than that for negative prediction errors (Fig. 1G). Comparing the pulse occurrence rate in the empirical data with the estimated learning rate, the learning rate for positive prediction errors was higher than that for negative prediction errors in the data with higher pulse occurrence rates during extinction (Fig. 3D). The linear mixed model revealed significant effects of the reward probability (*F*_(3, 79.00)_ = 4.45, *p* = 0.006), and the predicted pulse ratio was larger for the highest reward probability than for the lowest two reward probabilities (1.00 vs. 0.33, *t*_(70.53)_ = 2.73, *p* < 0.001; 1.00 vs. 0.25, *t*_(69.91)_ = 3.36, *p* = 0.007). Comparing the ratios between the positive and negative learning rates across conditions, the ratio increased as the reward probability increased (Fig. 3E; *t*_(81)_ = 11.33, *p* < 0.001). Furthermore, the higher were the weighted sum of behavioral value and curiosity during the reward phase estimated by the model, the higher was the response rate during the reward phase in the empirical data, indicating that Q-HCM successfully captured not only the pulse during extinction, but also behavioral features during the reward phase (Fig. 3F; *t*_(81)_ = 4.22, *p* < 0.001). These results imply that the model incorporating curiosity correctly captures the behavioral features of mouse lever pressing compared to the model that does not.

## Discussion

In this study, we revealed the psychological and computational mechanisms underlying an extinction burst. First, we built a curiosity-driven reinforcement learning model, Q-HCM (Fig. 1A). The model estimates the expected reward prediction error as curiosity by hierarchically calculating prediction errors and accumulating second-order prediction errors over time. Second, we conducted simulations to specify the determinant of extinction bursts, and revealed that reward probability controls the likelihood of extinction bursts (Fig. 1G). In addition, our model demonstrated a unique behavior, in that the higher response strength in the reward condition was uncertain (Fig. 1H). We also conducted computer simulations using the standard reinforcement learning model, but the burst did not occur with this model, suggesting that the curiosity-driven model is essential for the generation of extinction bursts. Third, we tested whether reward probability controlled the extinction burst using an operant conditioning task with head-fixed mice. The experimental results were consistent with the simulations, and Q-HCM fit the empirical data well (Figs. 2 and 3). These results provide theoretical and empirical evidence that the curiosity-driven model mediates extinction bursts.

As we concluded that curiosity mediated the burst, we also assumed that the burst is a type of information-seeking behavior (12, 13, 15, 42, 43) because *v* interpreted as curiosity reflects the information brought by the response. The information brought about by an event is defined mathematically as − *log* log *p*, where *p* denotes the probability of the event (44). Therefore, the smaller the probability is, the greater is the amount of information, implying that unpredictable events bring large amounts of information. The absolute value of the reward prediction error is similar to that of the information because it is larger when reward delivery is unexpected. The expected value of the information, information entropy, is *p* log *p* + (*1* − *p*) log(*1* − *p*) and the form of the equation is an inverted U-shaped curve, with a maximum at *p* = *0*.*5* and a minimum at *p* = *0* and *p* = *1*. The expected value of reward prediction error *v* has qualitatively the same form as the information entropy. In this sense, we call *v* curiosity, and the burst can be considered a type of information-seeking behavior.

Although extinction bursts are often mentioned in psychology textbooks (29, 45), there are few reports of experimental situations (32, 33). The two primary differences between existing studies and this study are the experimental system and reward probabilities. Existing studies on the effects of extinction on operant responses have been conducted using freely moving animals. Extinction induces a variety of behaviors, including aggression, social interaction, and drinking (34–36, 46), or increases the variability of operant responses (26). These experimental facts suggest that the secondary effects of extinction do not necessarily occur solely in one specific dimension of the operant response, that is, the response rate. By contrast, we used a head-fixed experimental setup for the mice. Because there is little behavior to engage in other than the operant response under the head-fixed situation, extinction bursts may be more likely to occur than under the free-moving experimental setup because the side effects of extinction on various behavioral dimensions do not occur. Our simulations with Q-HCM suggested that reward probability controls extinction bursts are sensitive to changes in reward probability (Fig. 1G). The behavioral results in mice also showed that bursts clearly occurred under conditions with the highest reward probability, but the bursts attenuated drastically as the reward probability decreased (Fig. 2D, E, G, and I). In contrast to our experiment, it is likely that extinction bursts did not occur in previous studies because the animals were trained at lower reward probabilities than in the present study (32, 33, 47) and there were a few cases reporting that extinction bursts occurred with an extremely high reward probability (48–50). Altogether, extinction bursts may likely occur in situations where animals have limited behavioral options in their repertoires and extremely high reward probability situations.

Our results suggest that extinction bursts are a type of information seeking. However, they may also be a complex phenomenon in natural situations. There are two alternative explanations for the extinction bursts. The response-competition theory attempts to explain extinction bursts by response competition between the operant response and experiment-unrelated behavior (47). Rewards could work not only to reinforce a specific response but also to induce other byproduct behaviors that are not directly related to the experiment (36, 51, 52). When reward events occur frequently, the operant response is disrupted by the induced behavior and the frequency of the operant response diminishes. Once extinction begins, the experiment-unrelated behavior induced by the reward vanishes, and the operant response increases transiently because of the less competitive behavior (47). According to this response-competition theory (47), neither the extinction burst nor suppression of responses under high reward probabilities in the reward phase are predicted to occur if competing responses are not induced by the reward. However, in our experiments, the use of a head-fixed apparatus limited the mice from engaging in experiment-unrelated behaviors. This experimental situation might be suitable for mice to show the extinction burst and decrease the response rate at higher reward probabilities (Fig. 2 D and G, H, J, and L). The fact that the extinction burst and downturn response rates were observed at higher reward probabilities in the head-fixed experimental setup suggests the involvement of curiosity in mouse behavior.

Another hypothesis that intends to explain extinction bursts is frustration theory. Frustration theory explains that extinction bursts are caused by the frustration that animals face after reward omission (53–56). There are difficulties in employing frustration theory to explain elimination bursts for three reasons. First, much of the evidence supporting frustration theory comes from experiments using double runways with running speed as the measure. Most extinction studies have used the frequency of key pecking by pigeons or lever pressing by rats in operant boxes as a measure; such experimental and measured differences limit the application of frustration theory in free-operant situations. Second, some studies have reported an increase in response frequency due to frustration following reward omission in the operant responses of pigeons and rats in the operant box (56, 57), but have been dismissed by response-competition and timing theories (47, 58). Another problem with frustration theory is that it does not have a rigorous mathematical definition, unlike the response-competition theory and our model. This undermines the scientific utility of frustration theory because it cannot be used to predict the results of a particular experimental situation or identify the variables that contribute to extinction bursts. Therefore, there is little room for frustration in explaining extinction bursts. However, it remains possible that frustration may also be involved in various phenomena associated with extinction. Animals display various behaviors after reward omission, such as aggression, emotional expressions, and escape behavior (36, 50, 59–61), which might be related to exploring behavior after the negative prediction error. Additionally, extinction increases the concentration of corticosterone in the blood (62). This increase in the corticosterone concentration began five minutes after the start of extinction and reached a maximum at 20 minutes. Further examination of the detailed temporal distribution of schedule-induced behaviors revealed that operant responses were observed for several tens of seconds immediately after the onset of extinction and aggressive behaviors followed the operant responses (36, 50). While these stress-related physiological measures and aggressive behaviors were observed in time windows ranging from tens of seconds to several minutes, the extinction burst in our study was observed in a very short time window of a few seconds (Fig. 2E, F, G, H, and L). The difference in the time windows suggests that extinction bursts and other extinction-induced behaviors may be mediated by different processes.

The behavioral characteristics of extinction bursts and their computational mechanisms are similar to those of obsessive-compulsive disorder (OCD). OCD is characterized by a compulsion to repeat certain behaviors or confirmations derived from obsession (63, 64). For example, one may wash one’s hands repeatedly because of the belief that one’s hands are dirty, even if one washes one’s hands repeatedly. From the standpoint of our model, extinction bursts are a type of information-seeking or checking behavior that confirms the omission of a reward by engaging in a repetitive operant response when the reward is unexpectedly omitted, similar to OCD. Furthermore, patients with OCD have been reported to have impairments in extinction learning (64–66), and their obsessive-compulsive behavior may be due to difficulties in extinction obsession. Our model predicts that elimination bursts are more likely to occur when the learning rate for negative reward prediction errors is smaller than that for positive reward prediction errors, and that individuals who actually experienced the extinction burst had smaller estimates of negative reward prediction errors (Fig. 1G and 3E). Thus, we suggest that the extinction burst is caused by persistent reward prediction due to slower extinction learning that has already been acquired, similar to the mechanism of obsessive-compulsive symptoms in OCD.

In conclusion, we provide theoretical and experimental evidence that operant responses in mice are directly controlled by external rewards and intrinsic factors such as curiosity. Although primary rewards powerfully drive animal behavior, intrinsic factors such as curiosity also play a role in modulating behavior. However, it is difficult to examine how intrinsic factors influence behavior compared to external rewards, which can be directly observed. To overcome this challenge, we implemented a curiosity-driven reinforcement learning model that predicts the emergence of curiosity and its influence on behavior. We predicted that phasic bursts of responses would occur immediately after extinction and extinction bursts would occur in trained responses with a high reward probability. This prediction was validated through behavioral experiments on mice. By combining computational modeling and behavioral experiments, we successfully evaluated the effects of intrinsic factors on behavior, which were previously difficult to evaluate. In the future, it is expected that this experimental system will be used to identify the psychological and neural mechanisms of curiosity by identifying the factors controlling extinction bursts and the brain regions and neural circuits behind these bursts.

## Materials and Methods

### Simulations

To identify the conditions under which the extinction burst occurred in Q-HCM, we calculated the intensity of the burst using various model parameters and parameters characterizing the environment. Burst intensity was defined as the increase from the asymptote of the response intensity under a given reward probability produced by extinction. We calculated the expected values of *Q* and *v* to determine the asymptote. Depending on whether a reward is presented, the reward prediction error (eq. 1) can be rewritten in the following two ways: δ^+^ = *1* − *Q* and δ^−^ = *0* − *Q*. The probabilities of positive and negative reward prediction errors depend on reward probability *p*; thus, eq. 2 can be rewritten as:

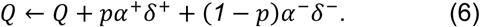

Similarly, we rewrite eq. 3 as follows:

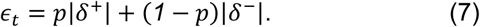

By updating *v* by redefining ϵ_*t*_ and eq. 4 and updating *Q* with eq. 6, we obtain the response intensity at a given time step *t* under arbitrary model parameters and reward probabilities. We calculate the response intensity with eqs. 4, 6, and 7 for a given reward probability until the change in the response strength per step becomes sufficiently small (<0.00001). After the response strength converged, we set the reward probability to zero and updated the response strength using eqs. 4, 6, and 7, until the response strength becomes less than that of the asymptote. The sum of the difference between the response strength and the asymptote from the onset of extinction until the response strength fell below the asymptote was considered as the intensity of the extinction burst.

### Subjects

Eight adult male C57BL/6J mice were used. All mice were purchased from Nippon Bio-Supp. Center and bred in a breeding room provided by the laboratory. All mice were naive and eight weeks old at the start of the experiment. The mice were maintained on a 12:12 h light cycle. All experiments were conducted during the dark phase of the light cycle. The mice had no access to water in their home cages and were provided with water only during the experimental sessions. The mice were allowed to consume a sufficient sucrose solution during the experiment. The mice’s body weight was monitored daily (22.84 ± 1.67 g before the experiment). They were provided additional access to water in their home cages if their body weight fell below 85% of the normal body weight measured before the experiment. Mice were allowed to feed freely in their cages. Experimental and housing protocols adhered to the Japanese National Regulations for Animal Welfare and were approved by the Animal Care and Use Committee of Keio University.

### Surgery

The mice were anesthetized with 1.0–2.5% isoflurane mixed with room air and placed in a stereotactic frame (942WOAE, David Kopf Instruments, Tujunga, CA, USA). A head post (H.E. Parmer Company, Nashville, TN, USA) was fixed at the surface of the skull, aligning the midline using dental cement (product #56849, 3M Company, Saint Paul, MN, USA) to head-fix the mice during the experiment. The mice were group-housed (four mice per cage) before the experiments, and a recovery time of 2 weeks was scheduled between the surgery and experiment commencement.

### Procedure

The mice were habituated to a head-fixed experimental setup (67–69) the day before the experiment commenced. During habituation, the mice were head-fixed in the apparatus under dim light and randomly presented with a 10% sucrose solution through a drinking steel spout. We presented a pure tone of 6,000 Hz at 80 dB during an experiment from a set of two speakers placed 30 cm in front of the platform and terminated the tone by the end of the experiment. After habituation, the mice were trained to press a lever placed in front of them. Mice could obtain a reward (10% sucrose solution) by pressing every lever, and the training was completed when the mice obtained 200 rewards in a session. If the mice stopped pressing the lever and did not reach 200 rewards in 1 h, the training session was completed and the mice were trained in the next session.

After the mice learned to press the lever, they were trained to press it with a specific reward probability. There were four reward probability conditions: 1.00, 0.50, 0.33, and 0.25. All mice experienced all conditions. Mice were trained under specific conditions for five sessions, and they could obtain 200 rewards in one session. After completing the training, a test session was conducted. In the test session, the mice obtained the reward by pressing the lever with a specific probability, but the extinction phase was inserted randomly four times within the session. The extinction phase continued until the mice stopped pressing the lever for 60 s, and the extinction phase finished; the mice could obtain the reward again until the next extinction phase. After the mice completed three test sessions, they were moved on to the next condition.

### Statistical Analysis

We collected data from all subjects repeatedly throughout our experiments and had missing values caused by sudden PC freezing for unknown reasons. We summarized the number of effective sessions per subject and condition (Table 1). Because our data were repeatedly measured and included missing data, we employed a linear-mixed model instead of a standard statistical analysis such as ANOVA. We set subjects and sessions as random effects throughout the analysis and present the fixed effects in the results section. We fitted a linear mixed model to our data using R (4.1 version) and the *lme4* package (70). We also used the *emmeans* package (71) to examine simple effects and employed Tukey’s method to adjust p-values for multiple comparisons.

**Table 1.** Number of effective sessions per subject and conditions.

| Subject ID | Reward probability for one lever pressing |  |  |  |
| --- | --- | --- | --- | --- |
|  | 1.00 | 0.50 | 0.33 | 0.25 |
| Sub 1 | 3/3 | 2/3 | 3/3 | 2/3 |
| Sub 2 | 3/3 | 3/3 | 3/3 | 3/3 |
| Sub 3 | 2/3 | 3/3 | 3/3 | 1/3 |
| Sub 4 | 3/3 | 3/3 | 2/3 | 2/3 |
| Sub 5 | 3/3 | 3/3 | 3/3 | 2/3 |
| Sub 6 | 3/3 | 3/3 | 2/3 | 3/3 |
| Sub 7 | 2/3 | 2/3 | 3/3 | 3/3 |
| Sub 8 | 2/3 | 3/3 | 3/3 | 2/3 |

Three test sessions were conducted for all conditions

### Pulse detection

We classified each response as a pulse (1) or not (0) based on the interval between one response and the previous response (inter-response time; IRT). First, we calculated reciprocal IRTs for all responses as local response rates and the 95% quantiles of the local response rates during the reward phase. We labeled a response as a pulse if its local response rate exceeded the 95% quantile. As the frequency of lever pressing varied by individual and day, this analysis was performed individually and by session.

### Model fitting to the empirical data

The Q-HCM and VQM were fitted to the dynamics of lever pressing in actual mice using Bayesian statistical modeling. For each subject’s session-by-session data, we estimated a set of parameters that would fit the pulse time series. We chose the pulse as the target variable for modeling because the heightened pulse rates corresponded to unusually fast response rates (i.e., extinction burst during the extinction phase). The main architecture of the model was almost the same as that of the simulation; however, only the behavioral output was different. As we fit the model to the binary time-series vector, including the given response as a pulse (i.e., {0, 0, 1, 0, 1 …0}), the model output changed from the simulation as follows:

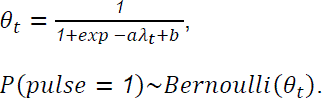

Here, λ_*t*_ is the response strength, defined as a weighted sum of action value and curiosity (eq. 5) and mapped into probability, λ_*t*_, using the logistic function. The pulse is generated from a Bernoulli distribution with parameter θ_*t*_. Hence, the pulse occurrence probability changes over time because the response strength, λ_*t*_, changes depending on agent actions and their outcomes. We fit the model to empirical data with the Bayesian inference framework Turing (72). The parameters and their prior distributions for Q-HCM and VQM are summarized in Table 2. All prior distributions were non-informative. We independently fitted both models to the data of each subject and session. We generated 4,000 MCMC samples for each model and data with 3,000 burn-in periods to discard the samples and four chains. Convergence was confirmed using the Gelman-Rubin statistic (R-hat) below 1.1. After the MCMC, we selected a better-fit model with widely applicable information criteria (73); WAIC). A no-U-turn sampler was used to generate the MCMC samples.

**Table 2.** Model parameters and prior distributions.

| Parameter | Prior distribution |  |
| --- | --- | --- |
|  | Q-HCM | VQM |
| $\alpha^+$ | *Beta*(1, 1) | *Beta*(1, 1) |
| $\alpha^-$ | *Beta*(1, 1) | *Beta*(1, 1) |
| $\alpha^v$ | *Beta*(1, 1) | - |
| $w$ | *Beta*(1, 1) | - |
| $a$ | truncated *Normal*(0, 100) | truncated *Normal*(0, 100) |
| $b$ | *Normal*(0, 100) | *Normal*(0, 100) |
| $q_0$ | *Beta*(1, 1) | *Beta*(1, 1) |
| $v_0$ | *Beta*(1, 1) | - |

## Data Availability Statement

The data supporting the findings of this study are available from the corresponding author upon request. The original code written for the analysis is available from the corresponding author upon reasonable request.

## Ethics Statement

The experimental and housing protocols adhered to the Japanese National Regulations for Animal Welfare and were approved by the Animal Care and Use Committee of Keio University.

## Author Contributions

KY and KT designed the study. KY and KT performed the stereotaxic surgery. KY collected all data from the head-fixed operant conditioning experiment with the help of KT. KY and HM conceptualized and built the computational model. KY analyzed all the data with the help of HM. KY and KT created figures. KY and KT wrote the manuscript, with comments from HM. KY, HM, and KT revised the manuscript.

## Funding

This research was supported by JSPS KAKENHI 18KK0070(KT), 19H05316 (KT), 19K03385 (KT), 19H01769 (KT), 20J21568 (KY), 22H01105 (KT), 23H02787 (KT), 23K27478 (KT), 23K22376 (KT), 24H00729 (KT), 24K16869 (KY), 24KJ0069 (KY), 24K16867 (HM), Keio Academic Development Fund (KT), Keio Gijuku Fukuzawa Memorial Fund for the Advancement of Education and Research (KT), and Smoking Research Foundation (KT).

## Conflict of interest

The authors declare that they have no conflict of interest.

## Acknowledgments

We would like to thank Shohei Kaneko, Ryuto Tamura, and Kazuko Hayashi for their assistance with animal care. We are also grateful to Kosuke Hamaguchi and Youcef Bouchekioua for the valuable discussions and for providing constructive comments on the manuscript.

## Notes

### Competing Interest Statement

The authors have declared no competing interest.

